# Dopamine release from Parkinson’s patient-derived neurons is disrupted due to impaired synaptic vesicle loading

**DOI:** 10.64898/2026.03.11.711038

**Authors:** Kaitlyn M L Cramb, Humaira Noor, Iona Thomas-Wright, Maria Claudia Caiazza, Sandor Szunyogh, Ira Milosevic, Dayne Beccano-Kelly, Stephanie J Cragg, Richard Wade-Martins

**Affiliations:** Oxford Parkinson’s Disease Centre and Department of Physiology, Anatomy & Genetics, University of Oxford, Oxford, OX1 3QU, UK; Kavli Institute of Nanoscience Discovery, University of Oxford, Dorothy Crowfoot Hodgkin Building, Oxford, OX1 3QU, UK; Aligning Science Across Parkinson’s (ASAP) Collaborative Research Network, Chevy Chase, MD, 20815, USA; Centre for Human Genetics, Nuffield Department of Medicine (NDM), University of Oxford, Roosevelt Drive, Oxford, OX3 7BN, UK; UK Dementia Research Institute at Cardiff University, Cardiff, CF24 4HQ, UK; School of Medicine, Cardiff University, Cardiff, UK

**Keywords:** Parkinson’s disease, dopamine release, alpha synuclein, iPSC, neurotransmission, synaptic vesicle, VMAT2

## Abstract

Striatal dopamine release defects are an early pathological feature observed in diverse models of Parkinson’s disease. However, the underlying molecular mechanisms responsible, and potential links to disease aetiology in humans, have been elusive. Here, we tested the hypothesis that dopamine release deficits are a characteristic feature of disease-relevant human neurons, using human Parkinson’s patient iPSC-derived dopamine neurons carrying the *SNCA*-triplication mutation. We reveal deficits in dopamine release from *SNCA*-triplication patient-derived neurons, and identify that this is due to reduced dopamine content arising from a lower capacity to store dopamine through reduced expression and function of vesicular monoamine transporter 2 (VMAT2) compared to healthy controls. In turn, by imaging VMAT substrate FFN206, and reporters for synaptic vesicular dynamics, SynaptopHluorin and CypHer5E, we reveal corresponding deficits in the size of either VMAT-containing, presynaptic releasing or recycling vesicle pools. Consistent with diminished synaptic vesicle loading and recycling, the cytosolic turnover of dopamine indicated by the ratio of concentrations of dopamine metabolite DOPAC to dopamine was elevated. By contrast, glutamate release events and VGLUT2 levels in neurons in the same preparations were not disturbed, demonstrating that vesicular dysfunction is limited to vesicles for dopamine. These findings therefore reveal dopamine loading into vesicles as a locus of dysfunction in human Parkinson’s-derived neurons. These disturbances will not only drive deficits in dopamine release but could potentially also be detrimental to dopamine neuron viability through an increased burden of oxidative stress associated with elevated cytosolic dopamine, thus contributing to both symptoms and aetiology of Parkinson’s pathology and offering a strategic target for improved therapies.

## Introduction

Deficits in dopamine release are a convergent hallmark of Parkinson’s disease models, typically presenting early, before dopamine neuron death and onset of motor symptoms, across a broad range of models of Parkinson’s disease.^1,2^ The mechanisms underlying dopamine release defects, and the potential impact of these mechanisms on the vulnerability of dopamine neurons to degeneration, are incompletely resolved. The protein α-synuclein, encoded by the *SNCA* gene, has several attributed roles in presynaptic function, but becomes a major constituent of Lewy bodies and is strongly associated with the pathology of Parkinson’s disease,^3^ neuropathologically and genetically. Abnormal levels of α-synuclein are implicated in familial and sporadic forms of Parkinson’s disease, with triplication of the *SNCA* locus resulting in a highly penetrant form of early-onset Parkinson’s disease.^4^ Corresponding rodent models of human α-synuclein overexpression, including the bacterial artificial chromosome (BAC)-transgenic (*SNCA*-OVX) mouse model that expresses wild-type human α-synuclein at elevated disease-relevant levels, or viral overexpression of human α-synuclein in rats, have identified deficits in dopamine release that can precede the progressive loss of dopamine neurons and motor dysfunction.^5–8^ These deficits in rodents have been linked to disturbances in synaptic vesicle biology, including alterations in vesicle density or clustering^5,6,8^ but the relevance of these phenotypes to disturbances in human neurons, and a direct mechanistic link to vulnerability to degeneration, has been undetermined.

Disruptions to vesicular biology will not only contribute to disturbances to neurotransmitter signalling, but might also promote vulnerability to degeneration.^9^ Diminished vesicular sequestration of dopamine can promote cytosolic metabolism of dopamine and cause dopamine to function as an endogenous neurotoxin.^10^ The loading of dopamine from the cytosol into synaptic vesicles is principally dependent on the activity of vesicular monoamine transporter 2 (VMAT2), an H^+^-ATPase antiporter encoded by *SLC18A2.*^11^ Knockdown of VMAT2 results in an accumulation of cytosolic dopamine, increased dopamine metabolism, dopamine neuron degeneration^12^ and age-related motor deficits^13–15^ comparable to behavioural deficits observed in rodent models of Parkinson’s disease, while in humans, the genetic missense mutation P387L in VMAT2 results in impaired dopamine loading and leads to an autosomal recessive severe infantile-onset movement disorder.^16^ Conversely, gain-of-function haplotypes in the *SLC18A2* promoter are associated with reduced risk of Parkinson’s disease,^17,18^ and enhancing VMAT2 levels in a BAC-transgenic mouse not only enhances dopamine release but also has the capacity to protect from the Parkinson’s-relevant neurotoxic insult MPTP.^19^ Thus, vesicular handling of dopamine, and by VMAT in particular, can serve dual purposes, supporting neurotransmission and limiting intracellular toxicity. However, few studies have addressed whether there is impaired dopamine loading into synaptic vesicles in human neurons in Parkinson’s disease.

Induced pluripotent stem cell (iPSC)-derived dopamine neurons (DANs) provide models for studying disturbances in cell biology and disease pathogenesis in human patient-derived cells. To better understand the mechanisms underlying dopamine release defects and connections to Parkinson’s disease aetiology, we characterised here the properties of cellular physiology, dopamine transmission, metabolism and handling in Parkinson’s patient-derived iPSC-DANs carrying *SNCA*-triplication. We report that while Parkinson’s iPSC-DANs with *SNCA*-triplication have similar physiological properties to human control DANs, by contrast, they exhibit reduced dopamine release, reduced synaptic vesicle pool size and disrupted dopamine loading into synaptic vesicles due to impaired VMAT2 function. This impairment will not only underpin dopamine transmission defects but also provide a key stepping stone towards an elevated metabolic burden.

## Materials and methods

Please see SI Appendix for comprehensive description of methods

### Cell culture

Human induced pluripotent stem cells (iPSCs) were handled as described on Protocols.io (DOI: dx.doi.org/10.17504/protocols.io.q26g7y1jqgwz/v1) and were differentiated into midbrain floor-plate derived DANs using a modified Krik’s protocol (20). Assays were performed at day 70 except when directly stated otherwise.

### Dopamine and glutamate release assays

KCl-evoked dopamine release was measured using high-performance liquid chromatography with electrochemical detection (HPLC-ECD). Briefly, dopamine release was evoked using 40 mM KCl in Ringer’s buffer (1.2 mM CaCl2, 40 mM KCl, 148 mM NaCl and 0.85 mM MgCl2, buffered to pH 7.4 using NaOH, 300 mOsm/kg) and total cell lysates were collected in 1% perchloric acid (PCA) for total dopamine cell content. Total cell concentration of monoamines was determined by HPLC-ECD using fresh mobile phase (13% methanol, 0.12 M NaH2PO4, 0.8 mM EDTA and 0.5 mM Octanesulfonic acid (OSA), pH 4.6). Samples were run on a 4.6 x 150 mm Microsorb C18 reverse-phase column and detected using Decade II ECD with a glassy carbon working electrode (Antec Layden) set at 0.7 V with respect to an Ag/AgCl reference electrode. Concentrations of monoamines were calculated compared to known standards. Evoked levels of glutamate release were measured using Glutamate Assay Kit (ab83389) according to the manufacturer’s protocol booklet.

### Whole-cell patch clamp electrophysiology

Whole-cell patch clamp of iPSC-DANs was performed using extracellular solution (167 mM NaCl, 2.4 mM KCl, 1 mM MgCl2, 10 mM glucose, 10 mM HEPES, 2 mM CaCl2, adjusted to a pH of 7.4, 300 mOsm at room temperature) and intracellular solution (140 mM K-Gluconate, 6 mM NaCl, 1 mM EGTA, 10 mM HEPES, 4 mM MgATP, 0.5 mM Na3GTP, adjusted to pH 7.3 and 290 mOsm). Series resistance (Rs) was maintained at <30 MΩ. Voltage gated channel currents were measured on voltage clamp where neurons were pre-pulsed for 250 ms with -140 mV pulse and a 10 mV-step voltage was applied from -70 mV to +70 mV.

### Live imaging assays

For the FFN206 assay, iPSC-DANs were incubated for 1 hour with 20 µM FFN206. When reserpine was used, cells were pre-treated for 30 minutes prior to assay. For the synaptotagmin-1-CypHer5E assay, Day 70 iPSC-DANs were treated overnight (Synaptic Systems 105 103CpH, 1:100) for 12 hours. Prior to imaging, cells were treated for 30 minutes with NeuroFluorTM NeuO (Stem Cell Technologies) and NucBlue (Thermo Fisher Scientific) as per manufacturers’ instructions. Images were subsequently taken on an Opera Phenix High-content screening system and spot properties were analysed using a custom pipeline developed with Harmony analysis software (v5.2, Revvity). For the synaptopHluorin live imaging, dopamine neurons were infected with Lentivirus on the final replating day (Day 20). Living imaging of synaptopHluorin experiments was done using a Perkin Elmer Spinning disk confocal and Nikon eclipse Ti microscope with a Hammatsu EM-CCD camera. Cells were imaged using 500 µL of classic Tyrode’s solution (124 mM NaCl, 1 mM KCl, 25 mM HEPES, 2 mM CaCl_2_, 1 mM MgCl_2_, 6 g/L glucose, pH 7.4), and when indicated, 500 µL of high stimulation Tyrode’s with 112 mM KCl (10 mM NaCl, 112 mM KCl, 25 mM HEPES, 4 mM CaCl_2_, 2 mM MgCl_2_, 6 g/L glucose, pH 7.4) was added as previously described ^20^, and maximum signal was measured by addition of ammonium Tyrode’s (69 mM NaCl, 2.5 mM KCl, 25 mM HEPES, 2 mM CaCl_2_, 2 mM MgCl_2_, 6 g/L glucose, 60 mM NH_4_Cl, pH 7.4) For details on fixed tissue immunocytochemistry, as well as protein and mRNA quantifications, please refer to supplemental information appendix.

### Statistical analysis

All statistical analyses were performed using GraphPad Prism 10.6.1 software. P-values < 0.05 were considered significant. An outlier test using Robust regression and Outlier removal (ROUT) method (Q=1%) was performed on all data. Details on specific statistical comparisons are outlined in figure legends.

## Results

### SNCA-triplication patient-derived iPSC-derived dopamine neurons have impaired dopamine release due to reduced dopamine content

Human iPSC-DANs generated from healthy controls and Parkinson’s iPSCs carrying the *SNCA*-triplication mutation (Figure 1A) were characterized using a range of techniques. iPSC-DANs expressed markers of dopamine neurons (Supplementary Fig. 1A), fired spontaneous and induced action potentials (Supplementary Fig. 1B, C), and released dopamine upon chemical stimulus (Supplementary Fig. 1D) in a calcium-dependent manner typical of neurotransmitter vesicular release (Supplementary Fig. 1E). iPSC-DANs carrying the *SNCA*-triplication mutation were confirmed to express 2-3-fold more α-synuclein than healthy controls (Supplementary Fig. 1F).

**Figure 1.**
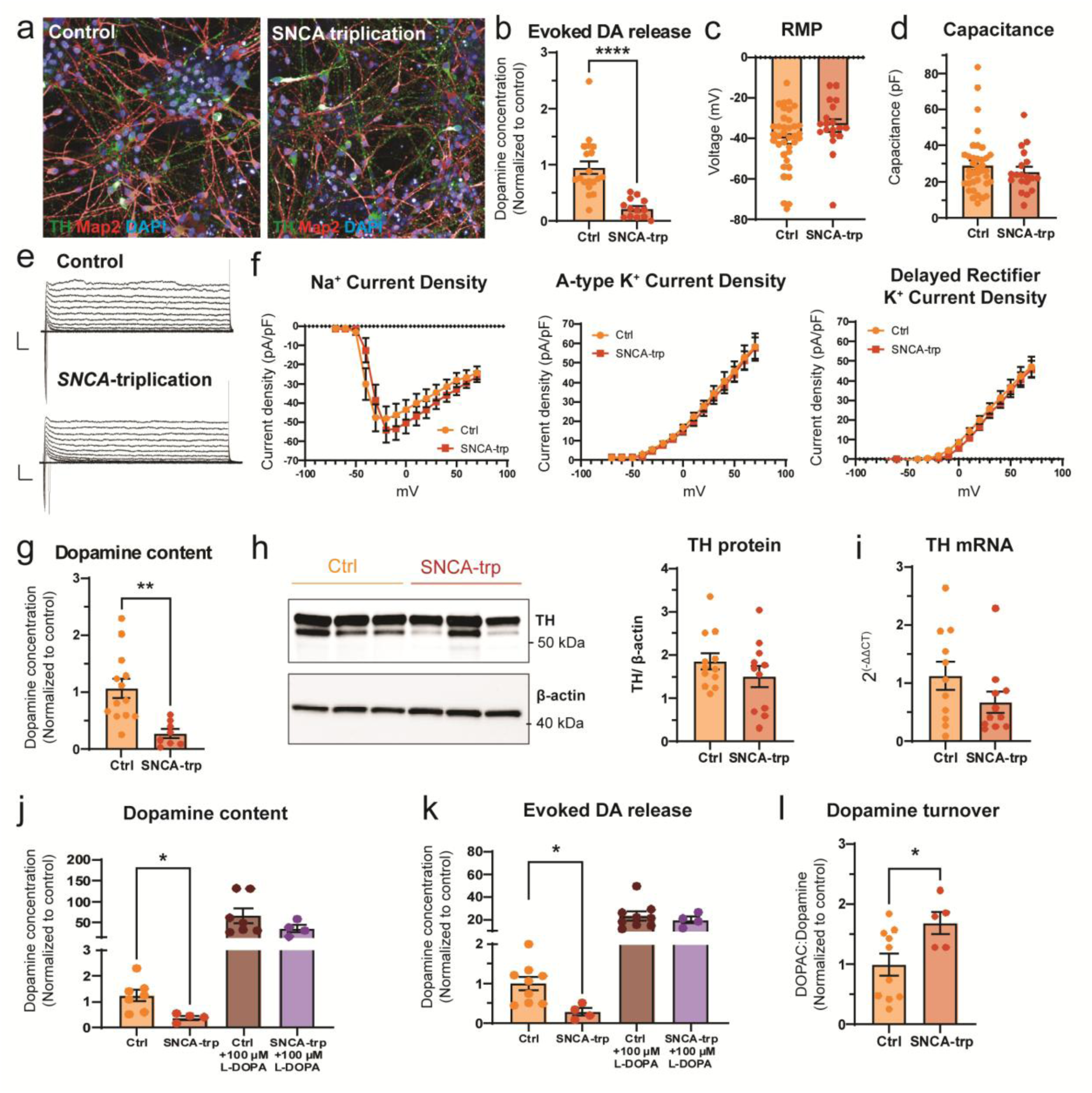
PD patient iPSC-derived dopaminergic neurons with SNCA-triplication have reduced dopamine release that is driven by reduced dopamine content independent of dopamine synthesis or maturation. (A) Example images of day 70 iPSC-DANs from healthy human controls and patients with the SNCA-triplication used in this study. (B) Dopamine release following chemical stimulation with 40 mM KCl as measured by HPLC-ECD. (C) Resting membrane potential and (D) Cell capacitance in voltage clamp mode. (E) Representative traces of whole cell patch clamp electrophysiology recordings in voltage clamp mode held at -70 mV with 400 ms voltage steps of 10 mV to +70 mV. Y-axis represent 500 pA and x-axis represents 20 ms. (F) Current densities of inward and outward currents as quantified by current normalized to capacitance. (G) Total cellular dopamine content as measured by HPLC-ECD. (H) TH protein expression by western blot and corresponding densitometric quantification. (I) RT-qPCR quantification of TH expression. (J) Dopamine content and (K) KCl-evoked dopamine release following 30 minute 100 µM L-DOPA treatment. (L) Dopamine turnover as measured by ratio of DOPAC to dopamine. Data points represent individual lines from 5 (B), 4 (H,I), 3 (G), or 2 (J,K,L) differentiations, or individual neurons from 5 control and 3 SNCA-triplication lines over 5 differentiations (C,D,E,F). Statistical analysis performed using an unpaired two-tailed Student’s t-test (B,C,F,G,H,I,L) and an ordinary one-way ANOVA with Tukey’s test for multiple comparisons (J,K); * denotes *p*-value <0.05, ** denotes *p*-value <0.01, *** denotes *p*-value <0.001. Ns = non-significant. Scale bar represents 50 µm (A).

Next, we investigated the ability of Parkinson’s patient neurons to release dopamine. Human patient-derived *SNCA*-triplication iPSC-DANs exhibited decreased dopamine release compared to control iPSC-DANs, when detected in extracellular medium using HPLC following a chemical stimulus of 40 mM KCl (Figure 1B). We tested whether the different release properties might be attributed to differences in neuronal excitability, but no differences were observed between control and *SNCA*-triplication iPSC-DANs in resting membrane potential or capacitance of these neurons (Figure 1C, D). We also measured inward and outward currents and observed equivalent densities of Na^+^ and A-type and delayed rectifier K^+^ channel currents when normalized to capacitance (Figure 1E, F). Therefore, dopamine release deficits are not driven by any gross differences in neuronal excitability, such as might be driven by differences in maturation. To confirm that the dopamine release deficit was not due to differences in the number of neuronal cells, resulting from differing differentiation efficiencies or cell death, we quantified the number of dopamine neurons. No differences in the number of dopamine neurons generated were observed between genotypes following differentiation at day 35 (Supplementary Fig. S2A, B) or day 70 when most assays were performed (Supplementary Fig. 2D, E), or in markers of cell death observed at either time point (Supplementary Fig. 2C, F).

We addressed whether diminished dopamine release resulted from a reduction in dopamine content and/or synthesis. Total cell dopamine content in *SNCA*-triplication iPSC-DANs was reduced compared to control iPSC-DANs (Figure 1G). However, expression levels of the main enzymes involved in dopamine synthesis were not different between genotypes for TH (protein, Figure 1H; mRNA, Figure 1I) or dopamine decarboxylase (DDC) (protein, Supplementary Fig. 3A; mRNA, Supplementary Fig. 3B). TH activity can also be altered by phosphorylation at Ser40 ^21,22^ and overexpression of α-synuclein has been shown to reduce phosphorylation at this site in MN9D and PC12 cells *in vitro*^22^ but p-Ser40TH levels were not different between *SNCA*-triplication and control iPSC-DANs (Supplementary Fig. 3C). Thus, while dopamine release deficits align with reduced content, these deficits are not likely to be due to underlying deficits in dopamine synthesis.

To assess whether deficits in dopamine release might be due to a limitation in dopamine storage machinery or capacity, we increased the driving force on dopamine storage by supplementation with a surplus of precursor L-DOPA. Treatment with 100 µM L-DOPA (as previously described^23^), substantially increased dopamine content (Figure 1J) and K^+^-evoked dopamine release (Figure 1K) and, furthermore, abolished the differences in both dopamine content and release between control and *SNCA*-triplication iPSC-DANs (Figure 1J, K). Lower levels of L-DOPA (20 µM) could be used to restore dopamine content and release in the *SNCA*-triplication iPSC-DANs to the levels seen in untreated control iPSC-DANs (Supplementary Fig. 3D-F), suggesting that dopamine release deficits arise from limited ability to accumulate dopamine when at endogenous levels.

If dopamine synthesis is normal, but vesicular uptake storage is compromised, we might expect this to be reflected in elevated cytosolic dopamine turnover owing to a prolonged presence in the cytosol and enhanced degradation by metabolic enzymes. Consistent with this hypothesis, we found that the ratio of the metabolite DOPAC to dopamine was greater in *SNCA*-triplication compared to control genotype (Figure 1L). This elevated ‘turnover’ could not be explained by any alterations in the expression of dopamine degradatory monoamine oxidases (MAO-A mRNA, Supplementary Fig. 3G, MAO-B mRNA and protein levels, Supplementary Fig. 3H, I). Taken together, these findings suggest that dopamine release defects are driven by an impairment in ability of neuronal cells to package endogenous dopamine, in turn leading to elevated cytosolic turnover.

### Dopamine loading into synaptic vesicles is impaired in SNCA-triplication patient-derived iPSC-DANs

We next tested the hypothesis that deficits in dopamine storage underpinning release are driven by impaired vesicle loading via VMAT2. We imaged fluorescent VMAT2 substrate FFN206^24^ and found fluorescent puncta were primarily localized to neuronal projections (Figure 2A) and the intensity of these puncta was decreased by the VMAT2 inhibitor, reserpine, in a concentration-dependent manner (Supplementary Fig. 4A). The number of spots was less in iPSC-DANs carrying the *SNCA*-triplication (Figure 2B), consistent with lower fluorescent spot intensity (Figure 2C), while the average area per spot was similar between genotypes (Figure 2D). Furthermore, we found that VMAT2 gene expression (*SLC18A2*) level was lower in *SNCA*-triplication iPSC-DANs (Figure 2E) suggesting that reduced dopamine release capacity results from compromised VMAT2-mediated vesicular uptake arising from more limited VMAT2 gene expression and VMAT2-expressing puncta. By contrast, mRNA and protein levels of the vesicular ATPase (v-ATPase) subunit VoA1 (*ATP6V0A1*) responsible for supporting the vesicular pH gradient required for VMAT2 function, were indistinguishable between genotypes (Figure 2F, Supplementary Fig. 4B).

**Figure 2.**
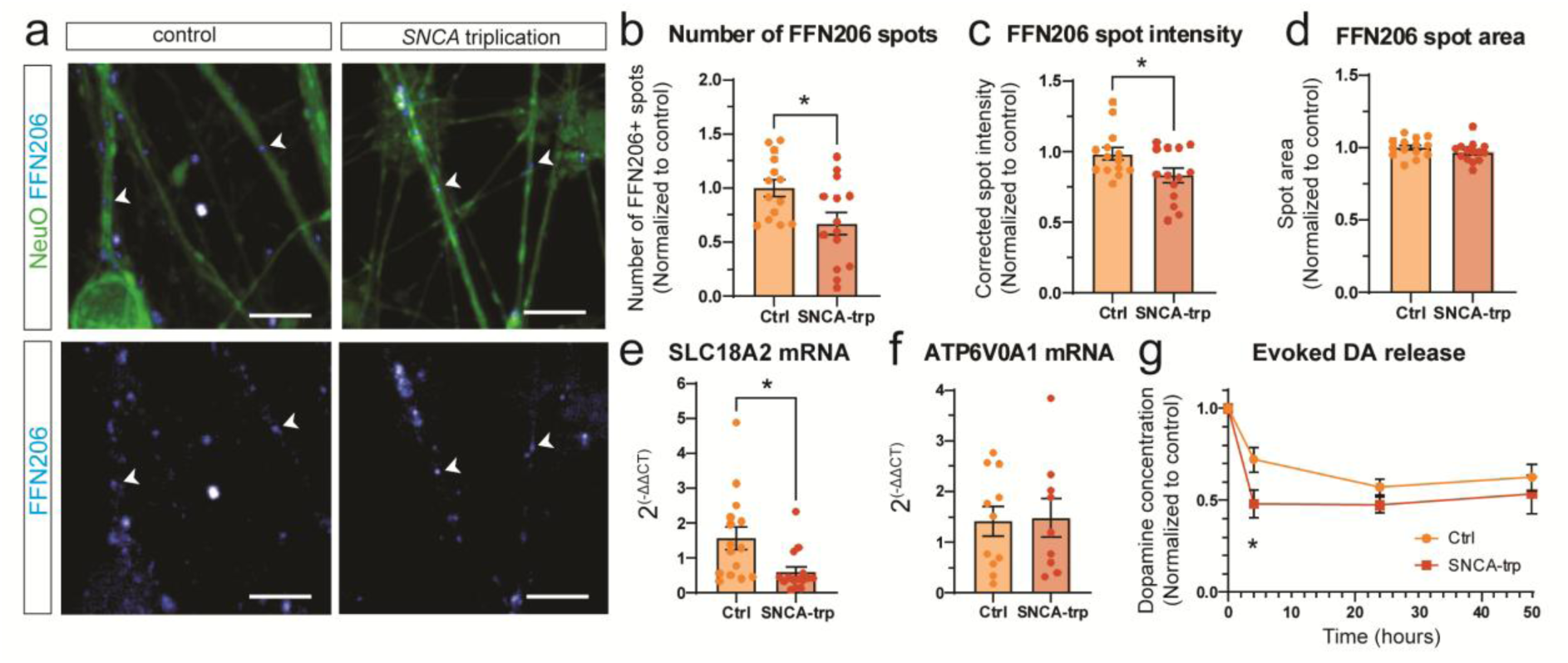
Dopamine loading in synaptic vesicles is impaired in PD patient-derived iPSC-DANs. **(A)** Example images of FFN206 uptake into iPSC-DANs. **(B)** The number, **(C)** corrected intensity and **(D)** area of FFN206 puncta as quantified from live imaging of FFN206. **(E)** RT-qPCR data of SLC18A2 mRNA encoding the protein VMAT2. **(F)** RT-qPCR data of ATP6V0A1 mRNA encoding the protein vATPaseA1. **(G)** Time course of dopamine release following repeated stimulations with 40 mM KCl. Statistical analysis performed using Student’s two-tailed t-test (B-F) or two-way ANOVA using Sidak test for multiple comparisons (G). * denotes *p*-value <0.05. ns = non-significant. Scale bar represents 10 µm. Data points represent individual lines over 5 (B-D), 4 (E) or 3 (F,G) differentiations.

To further corroborate the hypothesis that vesicle loading of dopamine is functionally impaired, we tested whether re-release of dopamine after an initial stimulus was more impaired in Parkinson’s than control cells. We quantified dopamine release at further timepoints (4, 24 or 48 hours) following an initial KCl-evoked release event and, when normalized to the level of dopamine released by the initial stimulation, dopamine re-release by a subsequent stimulation was less from iPSC-DANs carrying the *SNCA*-triplication than from control neurons (4-hours, Figure 2G), corroborating an impaired ability to re-load dopamine. Taken together, these data indicate there is disrupted dopamine loading into synaptic vesicles through a limited VMAT2 function.

### Synaptic vesicle pool size is disrupted in SNCA-triplication iPSC-DANs

We explored whether dopamine storage impairments associated with lower VMAT2 levels mapped on to a change in the readily releasable synaptic vesicle pool size. SynaptopHluorin, a fusion protein of synaptobrevin-2/VAMP-2 with a pH-sensitive fluorescent probe, allows the study of synaptic vesicle dynamics,^25^ as well as synaptic vesicle pool size,^26^ by acting as a vesicle release sensor that increases in fluorescence when vesicular pH becomes neutralized, e.g. in response to vesicle fusion with presynaptic membrane. We established expression of SynaptopHluorin in iPSC-DANs from healthy human controls (Figure 3A) using lentiviral transduction,^27^ confirmed that the application of NH_4_Cl to neutralize vesicles increased fluorescence intensity in localized varicosity-like regions along the axons (Figure 3B, Supplementary Fig. 4D), and confirmed that application of KCl to evoke vesicular exocytosis increased fluorescence intensity (Supplementary Fig. 4C,D). KCl increased fluorescence in only 63% of NH_4_Cl-responsive varicosities (Supplementary Fig. 4E), indicating that only a fraction of vesicle-containing varicosities are capable of responding to K^+^-evoked depolarisation. Approximately 30% of the maximum signal amplitude determined by NH_4_Cl was achieved after KCl stimulation (Supplementary Fig. 3F), identifying that within each responsive varicosity most vesicles in human dopamine axons are not immediately available for release following K^+^-evoked depolarisation. We then compared the level of NH_4_Cl-induced fluorescence between genotypes, and observed that the maximum NH_4_Cl fluorescence of individual varicosities was lower in *SNCA*-triplication iPSC-DANs (Figure 3C), consistent with a smaller total vesicle pool size.

**Figure 3.**
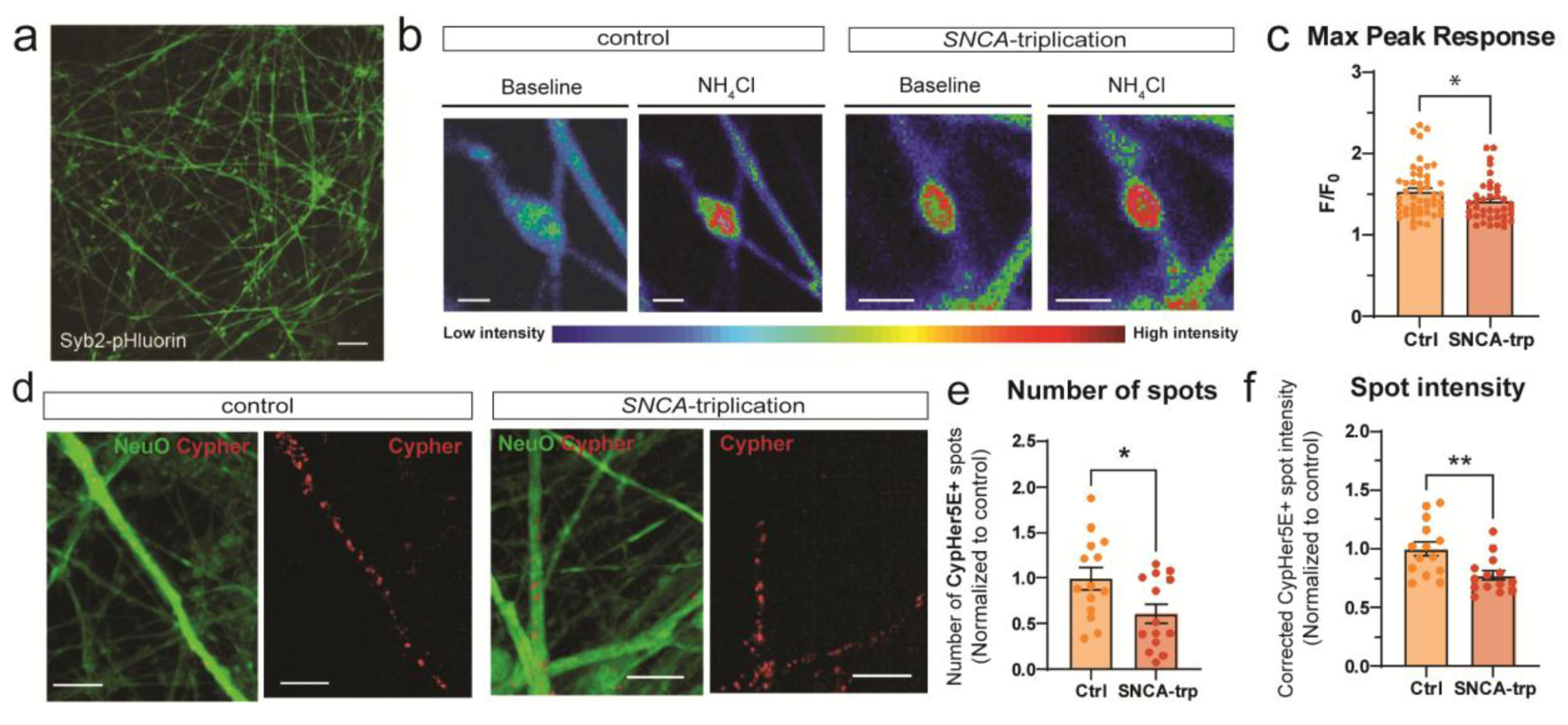
Impaired dopamine loading results in reduced vesicle pool size in PD patient-derived iPSC-DANs. **(A)** Representative image of day 70 iPSC-DANs expressing SynaptopHluorin. **(B)** Representative image of a varicosity at baseline (left), after and after addition of ammonium chloride (right) from control and SNCA-triplication iPSC-DANs. **(C)** Maximum fluorescence signal normalized to baseline intensity for healthy control and SNCA-triplication iPSC-DANs. **(D)** Representative images of healthy control and SNCA-triplication iPSC-DANs with the neuronal live marker (NeuO) and active synaptic vesicle marker (Synaptotagmin-1-CypHer5E). **(E)** The number and **(F)** corrected spot intensity of Cypher5E spots in iPSC-DANs. Dots represent individual varicosities (C) or individual lines from 4 differentiations (E,F). Statistical analysis performed using an unpaired Student’s two-tailed t-test (C,E,F). * denotes *p*-value <0.05, ** denotes *p*-value <0.01. Scale bar represents 50 µm (A) and 10 µm (B,D).

To confirm the smaller vesicle pool size in *SNCA*-triplication iPSC-DANs, we employed synaptotagmin-1-CypHer5E, a pH-sensitive variant of the Cy5 dye conjugated to an antibody against the luminal domain of synaptotagmin-1, that labels ‘active’ synaptic vesicles^28^ following its uptake by endocytosis as a part of actively recycling vesicles. A lower number of CypHer5E spots (Figure 3D, E) as well as fluorescent intensity of these labelled synaptic vesicles (Figure 3D, F) was observed in *SNCA*-triplication iPSC-derived neurons compared to controls. Notably, despite less total dopamine available for release, and a smaller synaptic vesicle pool size, we observed that *SNCA*-triplication iPSC-DANs have the same number of presynaptic and postsynaptic sites as well as the co-localization of the two (Supplementary Fig. 4G, H). These data suggest that *SNCA*-triplication iPSC-DANs do not compensate for the deficits in dopamine release and reduced synaptic vesicle pool size by increasing the number of synaptic sites.

### Glutamate release from SNCA-triplication patient-derived iPSC neurons is not impaired

Dopaminergic neurons are able to release other neurotransmitters, such as glutamate, and dysregulation of this co-release might contribute to the pathogenesis of Parkinson’s disease.^29,30^ While dopamine is loaded into vesicles via VMAT2, glutamate is loaded via VGLUT2. To assess whether vesicular loading deficits are specific to VMAT2 or generalizable to impact on VGLUTs, we investigated in *SNCA*-triplication iPSC-DANs whether glutamate transmission might also be impaired. VGLUT2, and not VGLUT1, was detected by RT-qPCR (Supplementary Fig. 1A), and both mRNA (Figure 4A) and protein (Supplementary Fig. 4I) levels of VGLUT2 were comparable between control and *SNCA*-triplication iPSC-DANs. Co-expression of VGLUT2 and TH in approximately 60% of neurons was confirmed using immunocytochemistry (Figure 4B), further confirming that glutamate can be stored by DANs. We note that these neuronal preparations do also contain non-TH+ neurons that might release glutamate making attribution of glutamate release characteristics to release from DANs alone difficult. Regardless, we observed no difference in the percentage of neurons positive for VGLUT2 (Figure 4B) and found that glutamate release was equivalent between control and *SNCA*-triplication iPSC-DANs when detected by ELISA (Figure 4C), or using whole cell patch clamp electrophysiology to record excitatory postsynaptic currents (EPSCs) in voltage clamp mode (Figure 4D). EPSCs (from local or recurrent inputs) were detected in approximately one third of neurons recorded in both genotypes (Figure 4E) with no differences in frequency or amplitude (Figure 4F, G). Combined, this suggests that the glutamate signalling in patient-derived iPSC-DANs from patients with *SNCA*-triplication is similar to healthy controls. Together, these data support a deficit in neurotransmission in human Parkinson’s patient-derived dopamine neurons that is specific to neurotransmitters dependent on VMAT2.

**Figure 4.**
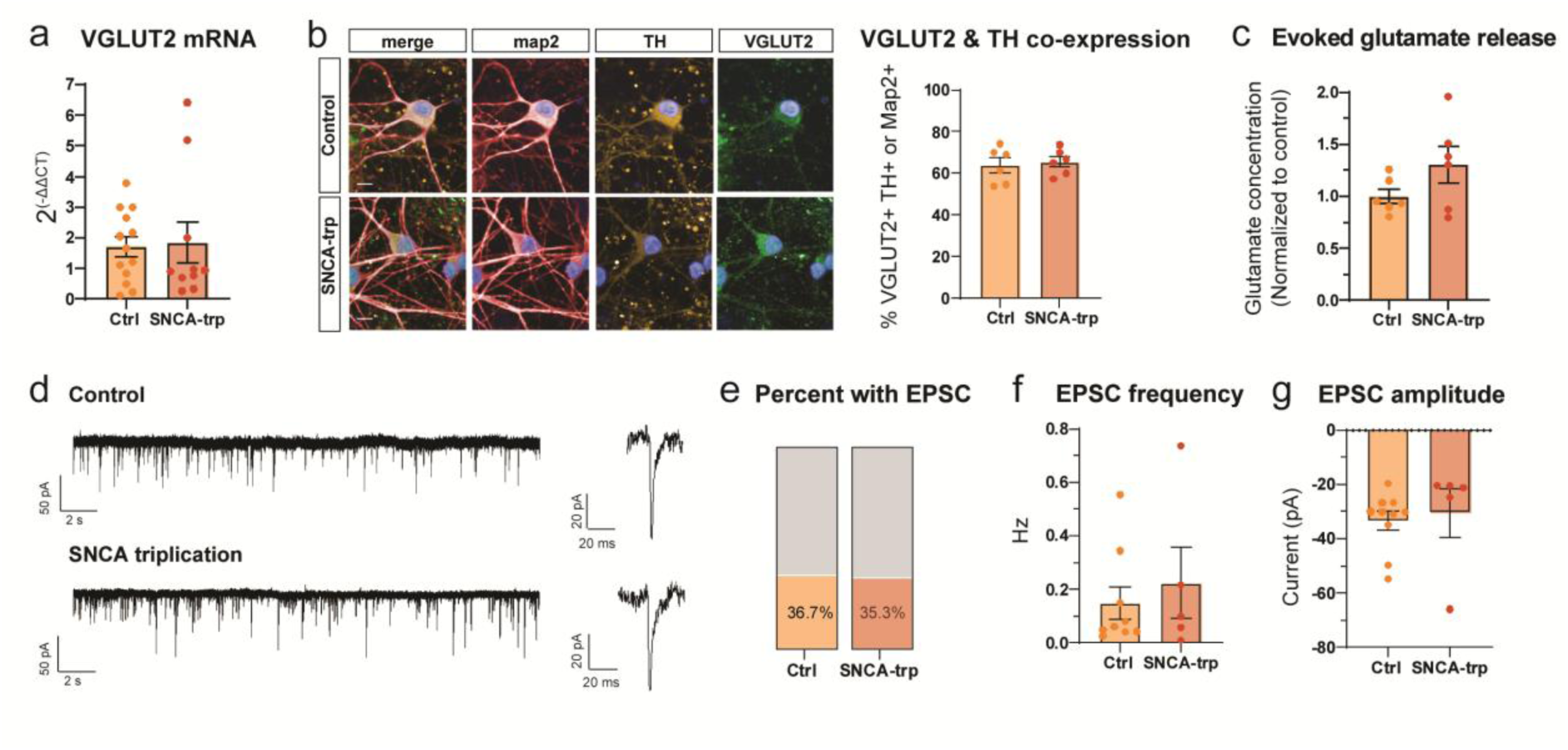
Glutamate release is unchanged in SNCA-triplication iPSC-DANs. **(A)** RT-qPCR quantification of VGLUT2 mRNA. **(B)** Representative images of iPSC-DANs expressing MAP2, TH and VGLUT2 and quantification of percentage of the total MAP2+ population also expressing VGLUT2 and TH. **(C)** ELISA quantification of 40 mM KCl-evoked glutamate release of day 70 iPSC-DANs. **(D)** Representative traces of day 70 iPSC-DANs on voltage clamp mode held at -70 mV. **(E)** Quantification of total number of cells with EPSCs. **(F)** The average frequency and **(G)** the average amplitude of the EPSCs. Scale bar represents 10 µm. Dots represent individual lines from separate differentiations (A,B,C) or individual neurons from 3 control or 3 SNCA-triplication iPSC-DANs from 5 differentiations (E,F,G). Statistical analysis performed using an unpaired two-tailed Student’s t-test. ns = non-significant.

## Discussion

Here, for the first time, we identify key characteristics of vesicular dopamine release from human iPSC-derived dopamine neurons, and reveal impairments in dopamine release from neurons derived from Parkinson’s patients. We identify underpinning deficits in vesicle pool size and VMAT2-dependent vesicle loading that are paralleled by corresponding elevations in cytosolic dopamine turnover. These vesicle loading deficits are attributable to deficits in VMAT2-expressing vesicles, and are not seen for VGLUT-expressing vesicles or glutamate release and handling in the same cultures, indicating a locus of dysfunction specific to VMAT2. Interventions aimed at enhancing vesicular loading of dopamine might therefore be a promising avenue for boosting dopamine transmission and supporting neuronal viability of remaining axons in Parkinson’s disease.

To our knowledge, these are the first observations to characterize fundamental properties of storage, release and turnover of dopamine vesicle pools in human iPSC-derived dopamine neurons. We find that dopamine release is limited by availability of precursor L-DOPA, involves VMAT2-dependent storage, and, as reported by imaging vesicle reporters synaptopHluorin, arises from only a subset of vesicle-containing varicosities, with some varicosities being functionally silent. These observations in human neurons match those in rodent striatum, where dopamine is released from only 20-30% of vesicle-containing varicosities at presynaptic active-zone like release sites^31,32^, indicating that functionally-silent, vesicle-containing axonal sites are a fundamental feature of dopaminergic neurons across species including human.

Furthermore, we provide evidence that the deficits in dopamine release seen in rodent models^5–8^ span across to human-derived iPSC dopamine neurons, therefore representing a unifying phenotype across parkinsonian burdens and species. Our finding that deficits in dopamine release from human iPSC-DANs derived from patients with *SNCA*-triplication arise from decreased VMAT2 function builds particularly on previous work suggesting impairments in dopamine loading in a variety of Parkinson’s disease models. Overexpression of α-synuclein or DJ-1 knockdown can reduce VMAT2 levels in SH-SY5Y cells ^33,34^ while VMAT2 expression is reduced in a *LRRK2*-G2019S mouse,^35^ and also altered in a *Vps35*-D620N knock-in mouse.^36^ We note further that a deficit in vesicular storage of dopamine does not extend to all neurotransmitters or vesicular transporters, as VGLUT function and glutamate transmission seems to be preserved, consistent with different neurotransmitters having distinct regulatory mechanisms and release properties even when co-transmitters.^37^

The implications of disrupted vesicular storage of dopamine in Parkinson’s disease could extend from maladaptive deficits in dopamine transmission to elevated disease burden. Although genetic variants in VMAT2 are not strongly associated with Parkinson’s disease in genome-wide association studies, rare familial mutations are known to cause an infantile-onset movement disorder similar to severe parkinsonism.^16^ VMAT2 inhibition by reserpine results in Parkinson’s-like behavioural symptoms^38^ while conversely increasing VMAT2 expression or function may be protective in dopaminergic neurons as increasing vesicular packaging not only rescues dopamine release defects but also protects against MPTP-induced neurodegeneration.^19^ Impairments in dopamine loading into synaptic vesicles not only reduces dopamine availability for release, but increases cytosolic dopamine whose turnover results in dopaminergic neuron degeneration through increased oxidative stress. We provide evidence to support the hypothesis that there is increased dopamine turnover in Parkinson’s patient-derived neurons, specifically an increase in the levels of dopamine metabolite DOPAC compared to dopamine. Increased dopamine turnover is observed very early in Parkinson’s disease using PET ^18^F-Fluorodopa imaging^39^ as well as in asymptomatic *LRRK2* mutation carriers.^40^ Appropriate metabolism and handling of dopamine is critical for maintaining neuronal health. When not stabilized by the acidic environment of a synaptic vesicle, dopamine is either enzymatically degraded or oxidized into neurotoxic species.^10^ Cytosolic dopamine has been suggested to be a major contributor to the vulnerability of dopamine neurons to degeneration in mice^41^ or Parkinson’s patient DANs with a range of Parkinson’s mutations, including *SNCA-*triplication.^42,43^ Chromaffin cells overexpressing Parkinson’s-variant *SNCA*-A30P also demonstrate increased cytosolic catecholamine levels.^44^ Dopamine catabolism includes the production of DOPAL catalysed by MAO and subsequent conversion to DOPAC catalysed by ALDH. DOPAL is particularly neurotoxic and is increased in post-mortem brains of Parkinson’s patients^45,46^ while genetic deletion of *Aldh* in mice reduces conversion to DOPAC and promotes dopamine neuron neurodegeneration.^47^ DOPAL can promote the formation of α-synuclein oligomers and cause damage to synaptic vesicles.^48^ Combined, the available evidence supports dopamine mishandling as a key component of Parkinson’s disease.

In this study, we provide critical corroboration in patient-derived neurons that dysfunction in dopamine neurotransmission prior to dopamine neuron death is a hallmark of Parkinson’s disease pathophysiology.^1,5,6^ We reveal that in *SNCA*-triplication patient derived neurons, impaired vesicle loading underlies a deficit in dopamine release, not shared by glutamate transmission. Given the potential deleterious ramifications of cytosolic turnover and toxicity of dopamine, strategies to enhance axonal vesicular loading of dopamine could offer a means to not only boost dopamine decaying transmission but provide a potential neuroprotective intervention of dual therapeutic benefit to Parkinson’s disease.

## Data availability

The data, protocols, and key lab materials used and generated in this study are listed in a Key Resource Table alongside their persistent identifiers at Zenodo (http://doi.org/10.5281/zenodo.14755599)

## Acknowledgements

This research was funded in part by Aligning Science Across Parkinson’s (ASAP-020370) through the Michael J. Fox Foundation for Parkinson’s Research (MJFF), in part by the Monument Trust Discovery Award from Parkinson’s UK (J-1403), and in part by a Wellcome Trust Collaboration Award (Grant ref. 223202/Z/21/Z), Wellcome Trust Investigator Award (224361/Z/21/Z) and La Caxia Foundation (HR22-00854). For the purpose of open access, the author has applied a CC BY 4.0 public copyright license to all author accepted manuscripts arising from this submission. KMLC is supported by a Junior Research Fellowship from St. John’s College and previously held studentships from the Natural Sciences and Engineering Research Council of Canada (PGSD3-517039-2018), the Canadian Centennial Scholarship Fund, St. John’s College and the Clarendon Fund.

## Supplementary material

Supplementary material is available online.

