## Supplementary for "Dopamine release from Parkinson’s patient-derived neurons is disrupted due to impaired synaptic vesicle loading"

#### This document includes:

Figures S1 to S4
Tables S1 to S4
Supplemental methodology
Supplemental references

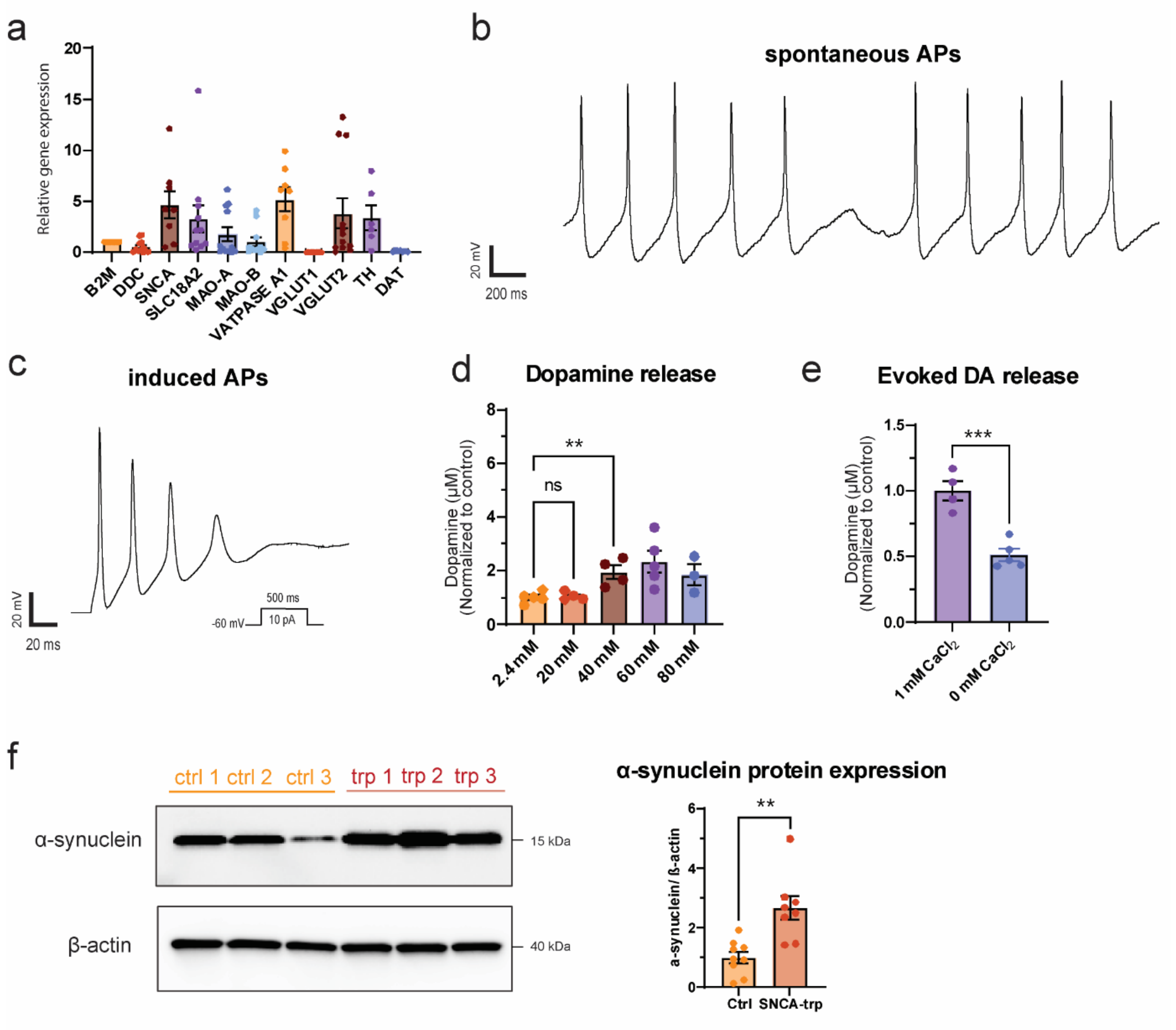


**Figure S1*****: Characterization of human iPSC-dopamine neurons used in this study***. (A) Relative gene expression of dopamine neuronal markers as quantified by qPCR. (B) Example traces of whole cell patch clamp electrophysiology demonstrating spontaneous action potentials at I=0 and (C) induced action potentials with iPSC-DANs held at -60 mV and current injected. (D) Dopamine release can be evoked with 40 mM KCl from day 70 iPSC-DANs. (E) Evoked dopamine release with 40 mM KCl is dependent on calcium. (F) Example Western blot and quantification from day 70 iPSC-DANs demonstrates α-synuclein protein expression is increased in day 70 iPSC-DANs with *SNCA*-triplication. For figures E-G dots represent individual healthy human iPSC lines from 3 (A,F) or 2 (D,E) differentiations. Statistical analysis performed using an ordinary one-way ANOVA with Tukey's test for multiple comparisons (D) or an unpaired two-tailed student's t-test (E,F). * denotes p-value <0.05, ** denotes p-value <0.01, *** denotes p-value <0.001. Ns = non-significant (p-value >0.05).

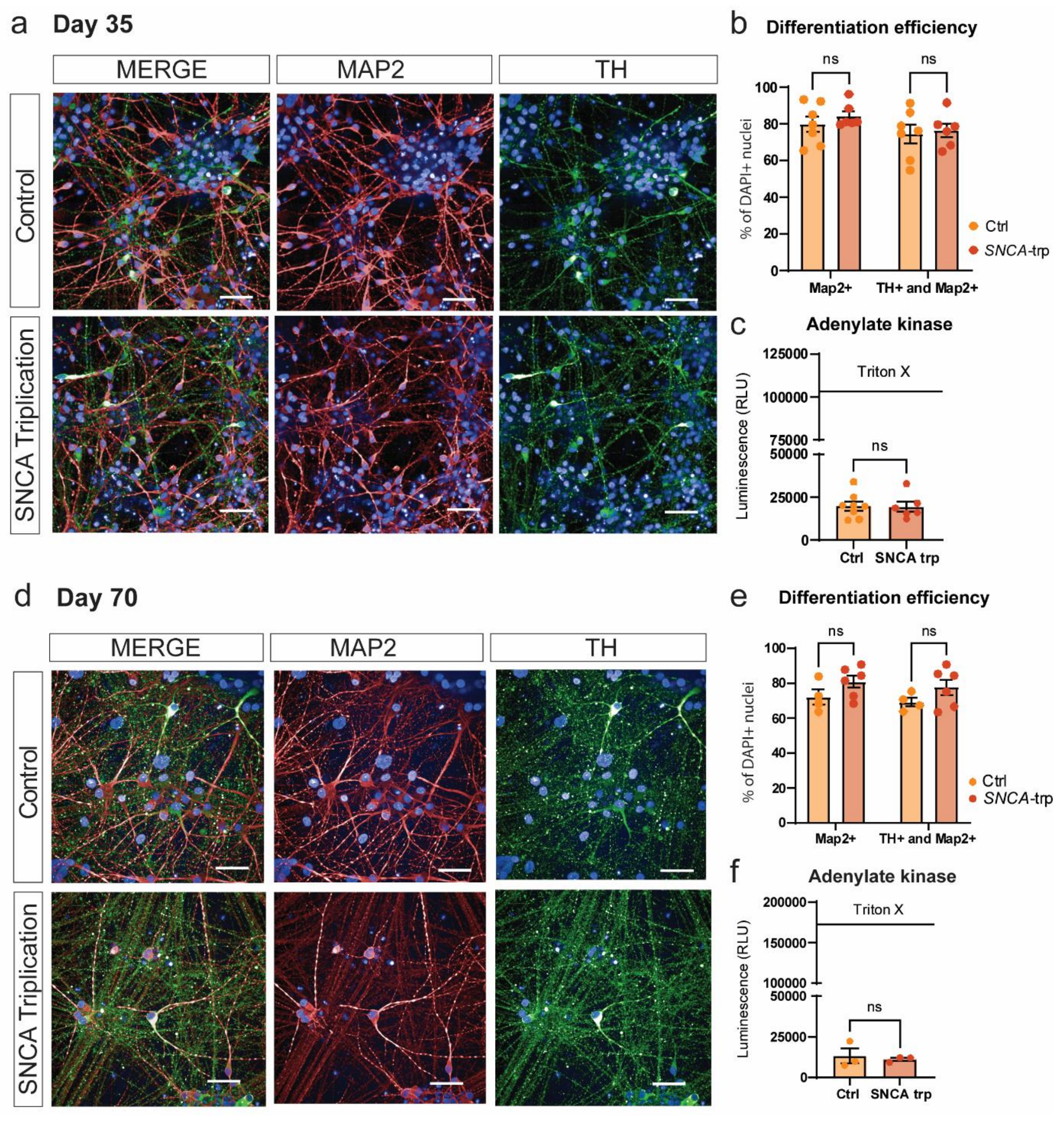


**Figure S2:** ***SNCA-triplication iPSC-DANs differentiate equivalently to those from healthy human controls and do not exhibit cell death phenotypes.*** Representative example immunocytochemistry images of day 35 neurons demonstrating differentiated cells express neuronal marker MAP2 and dopaminergic marker TH (A). These markers are expressed at equivalent levels between healthy and patient-derived iPSC-DANs with *SNCA*-triplication (B). No differences are present in the release of adenylate kinase between genotypes (C). Representative example images of day 70 neurons expressing TH and MAP2 at comparable levels across genotypes (D, E) and do not exhibit any differences in release of adenylate kinase, a marker of cell death (F). Scale bar represents 50 µm. n.s. represents non-significant (i.e. p-value >0.05). Statistical analysis represents an unpaired Student's two-tailed t-test (C,F). Data points represent individual lines from 2 differentiations (B,C,E) or 1 differentiation (F).

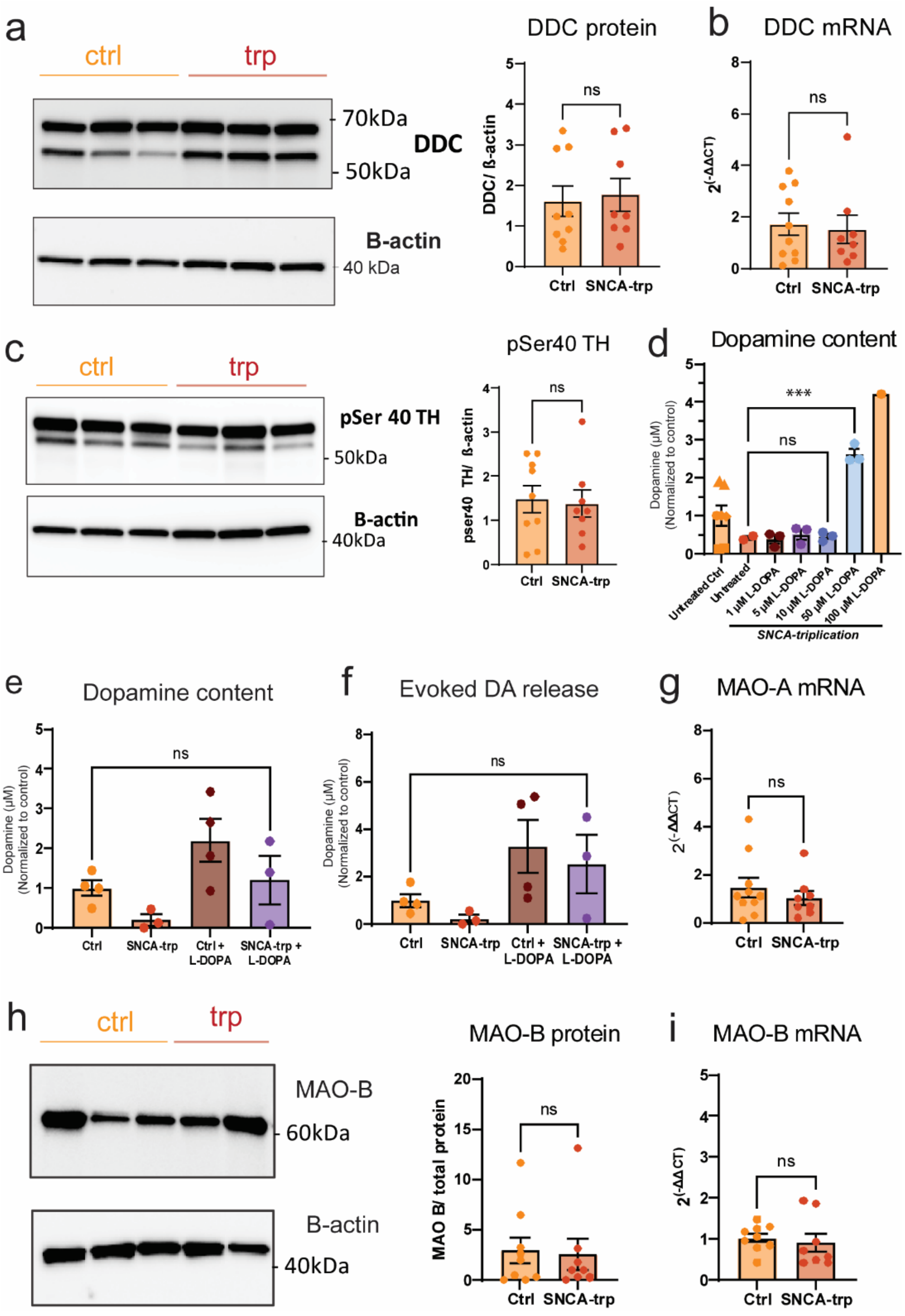


**Figure S3:** ***Dopamine synthesis and break down machinery is intact in iPSC-DANs with SNCA triplication and 20 µm L-DOPA rescue of dopamine content and release defects.*** (A) Representative example western blot for DDC in day 70 iPSC-DANs with *SNCA*-triplication or healthy controls and the densitometric quantification of band intensity normalized to β-actin. (B) mRNA expression of DDC as quantified by RT-qPCR. (C) Representative example western blot for phosphorylated TH and densitometric quantification of bands. (D) iPSC-DANs were treated with different concentrations of L-DOPA for 30 minutes, and the concentration needed to restore content levels of dopamine in iPSC-DANs with *SNCA*-triplication mutation to the level of untreated control was determined to be between 10 and 50 µM. (E) Dopamine content levels of iPSC-DANs with SNCA-*triplication* treated with 20µM L-DOPA for 30 minutes is equivalent to that of the untreated control. (F) KCl-evoked dopamine release of 20 µM L-DOPA treated iPSC-DANs with *SNCA*-triplication is equivalent to the untreated control. (G). RT-qPCR quantification of monoamine oxidase A (MOA-A) mRNA expression. (H) Representative example western blot of monoamine oxidase B (MOA-B) and densitometric quantification. Statistical analysis performed using an unpaired two-tailed Student's t-test (A,B,C,G,H,I) or an ordinary one-way ANOVA with Tukey's test for multiple comparisons (E,F). Data points represent individual cell lines from 3differentiations (A,B,C,G,H,I), 2 differentiations (D,E,F). *** denotes p-value <0.001. ns = non-significant (p-value >0.05).

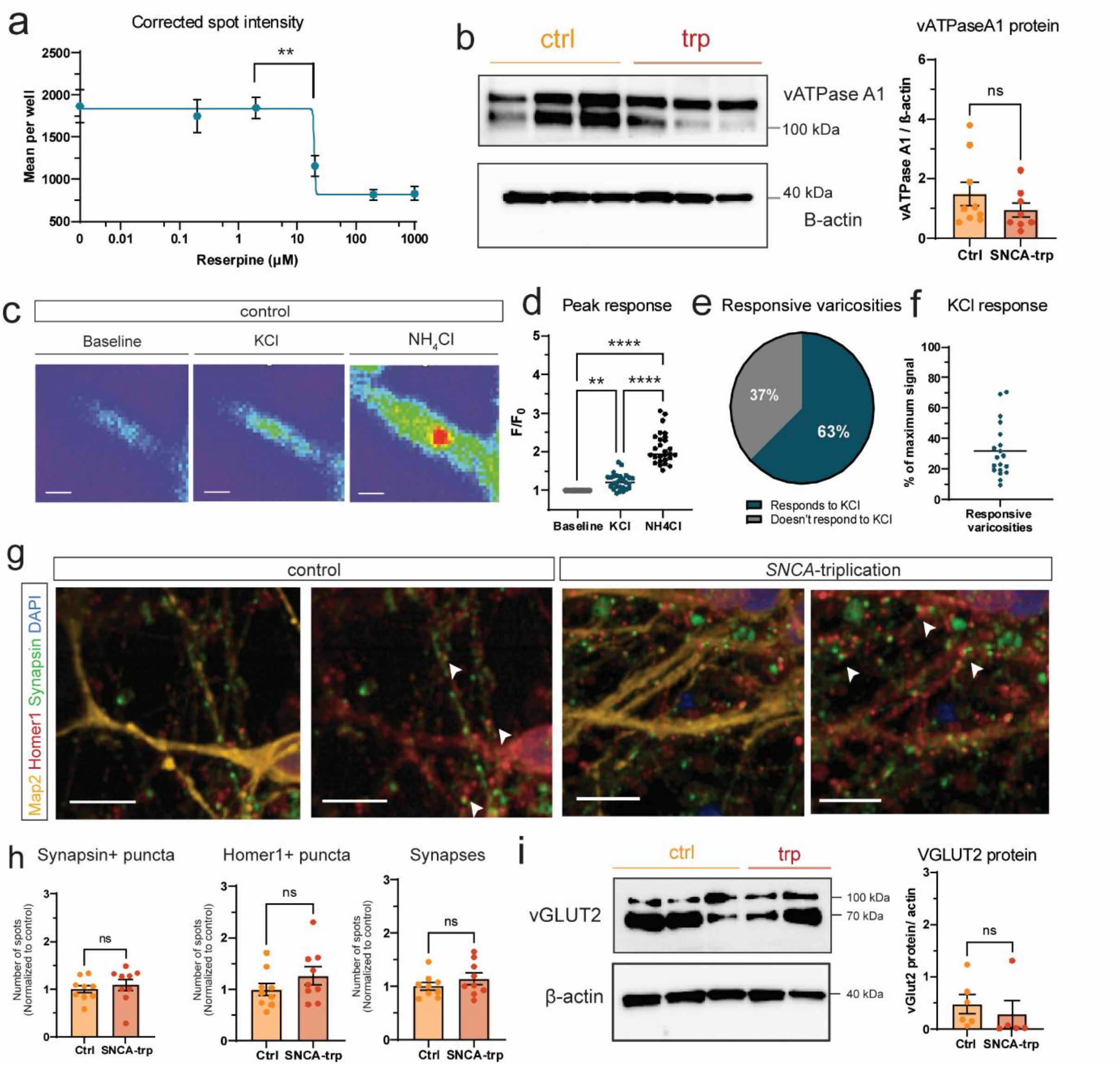


**Figure S4:** ***Physiological and synaptic properties in SNCA-triplication and healthy human control iPSC-DANs***. (A) Corrected spot intensity of FFN206-treated iPSC-DANs following treatment with reserpine. (B) Representative example western blot of vATPaseA1 protein and densitometric quantification. (C) Representative image of a varicosity at baseline (left), after high KCl stimulation (middle) and after addition of ammonium chloride (right). (D) Change of fluorescent intensity of varicosities relative to baseline after addition of high KCl or $NH_4Cl$; varicosities were determined by those that demonstrated an increase in intensity after $NH_4Cl$ and therefore contain synaptic vesicles. (E) Percentage of total $NH_4Cl$ responsive varicosities which show an increase in fluorescence of more than 10% ($F/F_0$ >1.1) after KCl stimulation. (F) Within individual varicosities of control dopaminergic neurons, the response to high KCl as a percentage of the maximum signal. (G) Representative example immunocytochemistry images of day 70 iPSC-DANs labelled with neuronal marker MAP2 or pre- and post-synaptic markers, Synapsin and Homer1, respectively. (H) Quantification of individual Synapsin+ puncta, Homer1+ puncta and co-localized Homer1+ and Synapsin+ puncta. (I) Western blot for VGLUT2 and densitometric quantification of total VGLUT2 protein normalized to β-actin. scale bars represent 10 μm. Dots represent individual varicosities (D,F) or individual lines from 3 differentiations (B,H,I). Statistical analysis performed using an ordinary one-way ANOVA with Tukey's test for multiple comparisons (D) or an unpaired Student's two-tailed t-test (B,H,I).* denotes p-value <0.05, ** denotes p-value <0.01, *** denotes p-value <0.001, **** denotes p-value <0.0001. ns = non-significant.

| 1º antibodies | Supplier | Catalogue # | Species | Dilution factor | Technique |
|---|---|---|---|---|---|
| Foxa2 | Abcam | ab108422 | Rabbit | 1:500 | ICC |
| Homer I | Synaptic systems | 160 003 | Rabbit | 1:500 | ICC |
| Synapsin I/II | Synaptic systems | 106004 | Guinea Pig | 1:500 | ICC |
| Map2 | Abcam | ab92434 | Chicken | 1:500 | ICC |
| TH | Sigma-Aldrich | ab1542 | Sheep | 1:500 | ICC |
| VGLUT2 | Synaptic systems | 135403 | Rabbit | 1:500 | ICC |
| VMAT2 | Thermo Fisher | PA522864 | Rabbit | 1:500 | ICC |
| $\alpha$-synuclein | Abcam | ab138501 | Rabbit | 1:10 000 | WB |
| DDC | Sigma | AB1569 | Rabbit | 1:1000 | WB |
| MAO-B | Abcam | Ab137778 | Rabbit | 1:1000 | WB |
| Phosphoserine TH | Sigma-Aldrich | AB5935 | Rabbit | 1:1000 | WB |
| TH | Millipore | AB152 | Rabbit | 1:1000 | WB |
| VMAT2 | Abcam | Ab70808 | Rabbit | 1:1000 | WB |
| vATPase (ATP6V0A1) | Abcam | Ab179858 | Rabbit | 1:1000 | WB |
| VGLUT2 | Synaptic systems | 135403 | Rabbit | 1:1000 | WB |

**Table S1:** Primary antibodies used in this study

| Secondary antibodies | Catalogue # (Supplier) | Dilution factor |
|---|---|---|
| Donkey anti-mouse IgG Alexa Fluor 488 | A21202 | 1:1000 |
| Donkey anti-chicken IgG Alexa Fluor 555 | A78949 | 1:1000 |
| Donkey anti-rabbit IgG Alexa Fluor 647 | A31573 | 1:1000 |
| Goat anti-guinea pig IgG Alexa Fluor 488 | Ab96954 | 1:1000 |
| Goat anti-chicken IgG Alexa Fluor 555 | A32932 | 1:1000 |
| Goat anti-rabbit IgG Alexa Fluor 647 | A21245 | 1:1000 |
| **DYE-** DAPI (4',6-Diamidino-2-Phenylindole, Dilactate) | D1306 | 1:10000 |

**Table S2:** Secondary antibodies used in this study

| Line Name | Genotype | Reprogramming method | Age/Sex | Reference |
|---|---|---|---|---|
| SFC840 | Healthy control | Cytotune (SeV) | 67F | (1) |
| SFC856-03-04 | Healthy control | Cytotune (SeV) | 78F | (2) |
| SFC065 | Healthy control | Cytotune 2 (SeV) | 65M | (3) |
| SFC067 | Healthy control | Cytotune 2 (SeV) | 72M | (4) |
| SFC156 | Healthy control | Cytotune 2 (SeV) | 75M | (4) |
| SFCAD2-01 (SBAd2-01) | Healthy control | Cytotune 1 | 51M | |
| SFC831-03-01 | SNCA triplication | Cytotune 1 | 55F | (2) |
| SFC831-03-03 | SNCA triplication | Cytotune 1 | 55F | (2) |
| SFC831-03-05 | SNCA triplication | Cytotune 1 | 55F | (2) |

**Table S3:** iPSC lines used in this study

| **Primer name** | **Sequence** |
|---|---|
| MAO-A Forward | GAG ACG CTG AAC CAT GAA CA |
| MAO-A Reverse | GGA AGC CGC TGA ATT AAC TG |
| MAO-B Forward | TTG GAG GAC AAT GGA TGA CA |
| MAO-B Reverse | CCA GCA GAG CTT GTC CAG TA |
| DDC Forward | AGG AAG CCC TGG AGA GAG AC |
| DDC Reverse | CAA AGG AGC AGC ATG TTG TG |
| ATP6V0A1 Forward | GGA TCC TTT CCC AGA AGA ATG |
| ATP6V0A1 Reverse | CAT GGA GAA CAC ACC CAT CA |
| VGLUT1 Forward | CAT GAG TGG TCT GGG CTT CT |
| VGLUT1 Reverse | TCT CTG GAT CCC AGC TGA AC |
| VGLUT2 Forward | TGT CAT GGG ATA TGG AGC AA |
| VGLUT2 Reverse | CTG CAC AAG AAT GCC AGC TA |
| TH Forward | CGA GCT GTG AAG GTG TTT GA |
| TH Reverse | CAC GAA GTA CTC CAG GTG G |
| VMAT2 Forward | CAG CTC ATC ACC AAC CCT TT |
| VMAT2 Reverse | AAG GCA TAG CTG CTG GAG AA |
| SNCA Forward | ATG TTG GAG GAG CAG TGG TG |
| SNCA Reverse | AAT TCC TTC CTG TGG GGC TC |
| B2M Forward | TTCTGGCCTGGAGGCTATC |
| B2M Reverse | TCAGGCAATTTGACTTTCCATTC |
| GAPDH forward | TCA TCA TCT CTG CCC CCT CT |
| GAPDH reverse | TCA TGG ATG ACC TTG GCC AG |

**Table S4:** qPCR primers used in this study

### Supplemental methodology:

#### Cell culture

Human induced pluripotent stem cells (iPSCs) were handled as described on Protocols.io (DOI: dx.doi.org/10.17504/protocols.io.36wgqjew3vk5/v1), plated on matrigel and expanded in mTeSR1 medium with mTeSR1 supplement (StemCell Technologies) and 100 U/ml Penicillin and 100 µg/ml Streptomycin (Life Technologies). Cells were passaged using EDTA or TrypLE (Life Technologies), replating with 10 µM ROCK inhibitor (Y-27632) (Tocris Bioscience). iPSCs were differentiated into midbrain floor-plate derived DANs using a modified Krik's protocol (5) as described on Protocols.io (DOI: dx.doi.org/10.17504/protocols.io.q26g7y1jqgwz/v1). Briefly, iPSCs were plated at 1.45 million/well on Geltrex into a well of a 6 well plate. Cells were passaged and replated at 1:1 once on day 9 using Accutase (StemCell Technologies) and expanded for 1 to 21 days in day 10 media. Cells were treated for 1 hour with mitomycin C on day 22 and subsequently fed 2 times a week with maturation media supplemented with antibiotic-antimycotic (Thermofisher). Cortical rat astrocytes (Life technologies, N7745100) were maintained in high glucose DMEM with 15%, 1x Pen/strep and 4 mM L-glutamine, fed every 4 days to a maximum of P3.

#### Dopamine release and content assays

KCl-evoked dopamine release was measured using HPLC-ECD as described on Protocols.io (DOI: dx.doi.org/10.17504/protocols.io.q26g7p2bkgwz/v1). Ringer's buffer (1.2 mM $CaCl_2$, 40 mM KCl, 148 mM NaCl and 0.85 mM $MgCl_2$ and buffered to pH 7.4 using NaOH and <300 Osm) was added to 500 000 cells/ well of a 24 well plate day 70 iPSC-DANs for 5 minutes following a single PBS wash. Supernatant was collected and placed in an Eppendorf containing 1 µL of undiluted perchloric acid (PCA). Samples were snap frozen on dry ice and stored at -80°C. Prior to loading on the HPLC column, samples were spun at 10 000 g for 10 minutes. The supernatant was then aspirated and pellets snap frozen on dry ice. Total cell concentration of monoamines was determined by HPLC-ECD. Total cell content was measured by collecting cell lysates harvested in 500 µL of 1% PCA. Prior to loading on column, samples were sonicated 10 times for 1 second and centrifuged at 10 000 g for 10 minutes. Fresh mobile phase (13% HPLC grade methanol, 0.12 M Sodium phosphate monobasic dihydrate ($NaH_2PO_4$), 0.8 mM Ethylenediaminetetraacetic acid (EDTA) and 0.5 mM 1-Octanesulphonic acid sodium salt (OSA), pH 4.6 and filter sterilised) was run at a flow rate of 1 mL/min. Samples were run on a 4.6 x 150 mm Microsorb C18 reverse-phase column and detected using Decade II ECD with a glassy carbon working electrode

(Antec Layden) set at 0.7 V with respect to an Ag/AgCl reference electrode. Concentrations of monoamines were calculated compared to known standards.

#### Whole-cell patch clamp electrophysiology

Day 70 iPSC-DANs plated at 100 000 cells/coverslip on 13 mm glass coverslips were placed in extracellular solution (167 mM NaCl, 2.4 mM KCl, 1 mM $MgCl_2$, 10 mM glucose, 10 mM HEPES, 2 mM $CaCl_2$ adjusted to a pH of 7.4 and 300 mOsm at room temperature). Electrodes with tip resistances between 7 and 12 MΩ were produced from borosilicate glass (0.86 mm inner diameter; 1.5 mm outer diameter). Intracellular solution (140 mM K-Gluconate, 6 mM NaCl, 1 mM EGTA, 10 mM HEPES, 4 mM MgATP, 0.5 mM $Na_3GTP$, adjusted to pH 7.3 and 290 mOsm) was added to the electrode. Data acquisition was performed using a Multiclamp 700B amplifier, digidata 1550A and clampEx 6 software. Series resistance (Rs) was maintained at <30 MΩ. Voltage gated channel currents were measured in voltage clamp mode where neurons were pre-pulsed for 250 ms with -140 mV pulse and a 10 mV-step voltage was applied from -70 mV to +70 mV. Induced action potentials were recorded in current clamp mode, neurons were held at -70 mV and 15 current steps of 10 pA were applied from -10 pA to 130 pA. Analysis was performed using Clampfit 10.3 (pCLAMP Software suite, Molecular Devices).

#### mRNA expression analysis using qRT-PCR

Dopamine neurons plated at a density of 500 000 per well of 24 well plate were collected using ice cold PBS, centrifuged for 5 minutes at 1000 xg and pellets were snap frozen on dry ice. Pellets were stored at -80ºC. RNA extraction was performed using the RNeasy Mini kit (Qiagen, 74104) according to the manufacturer's instructions and incorporating optional DNase step for on-column digestion of genomic DNA. Prior to elution, RNase-free $H_2O$ was prewarmed and incubated on column for 5 minutes to increase yield. Concentration and purity of RNA were determined using DeNovix DS-11 Series Nanodrop. cDNA synthesis was performed using Superscript III Reverse Transcriptase (Invitrogen). 400 ng of RNA was incubated with random primers and 10 µM dNTPs and incubated at 65ºC for 5 minutes. Next, 5x First strand buffer, 0.1 M DTT, RNase OUT and Superscript III Reverse Transcriptase were added. Samples were then incubated for 5 minutes at 25ºC, 50ºC for 60 minutes and 70ºC for 15 minutes. cDNA was stored at -20ºC. qPCR was performed using Fast SYBR Green Real-Time PCR Master Mix (Thermofisher) and a StepOnePlus Real-Time PCR System (Applied biosystems, Waltham, MA, USA). cDNA was diluted according to primers used, usually 1:20. Primers were added at 10 µM.

#### Western Blot

Neurons were harvest in PBS and lysed at 4°C in RIPA buffer (10 mM Tris-HCl, 1 mM EDTA, 1 mM EGTA, 140 mM NaCl, 1% Triton X-100, 0.1% SDS, 0.1% Sodium deoxycholate pH 7.4) with protease inhibitor (cOmplete Mini, EDTA-free, Roche). Protein concentration was determined by BCA assay (Pierce) according to manufacturer's instructions and adjusted to load 10 µg or 20 µg of protein per lane, in RIPA buffer. Samples were heated to 75°C for 10 minutes and resolved on Stain-free Midi-PROTEAN Precast Gels (Bio-Rad) for 60 minutes at 200V. Resolved proteins were imaged for total protein using the stain-free diazo dye and transferred to PVDF membranes using the Trans-Blot Turbo Transfer System (BioRad). Membranes were washed once in tris-buffered saline containing 0.1% Tween-20 (TBS-T) and fixed for 30 minutes in 4% formaldehyde (ThermoFisher Scientific) diluted in Milli-Q water (Merck) in order to reduce loss of small proteins as previously described (6). Membranes were washed >10 times with Milli-Q water and then blocked for 1 hour in 5% skimmed milk in TBS-T. Blots were incubated overnight at 4°C with primary antibodies (Table S1) in 5% skimmed milk in TBS-T. Blots were washed with TBS-T 3 times for 10 minutes with agitation, and incubated with secondary antibodies for 1 hour at room temperature. Detection was performed with Amersham ECL substrate and quantified with BioRad ImageLab.

#### Glutamate release assay

Day 70 iPSC-DANs plated at 250 000 cells/well of 12 well plate were washed once with phosphate buffered saline (PBS). For evoked release quantification, 200 µL of prewarmed HBSS++ (with 40 mM KCl) was added to the same wells and incubated for 5 min. Conditioned media was collected in both instances and stored at −20 °C until further use. Evoked levels of glutamate release were measured using Glutamate Assay Kit (ab83389) according to the manufacturer's protocol booklet. Protein concentration of the samples was quantified using Pierce™ BCA Protein Assay Kit (ThermoFisher) following the manufacturer's instructions. The OD was read at 450 nm in a PHERAstar® microplate reader (BMG Labtech). Protein concentration and glutamate concentrations were determined from their respective standard curves. Detailed protocol can be found at Protocols.io (DOI: dx.doi.org/10.17504/protocols.io.j8nlkoqdxv5r/v1).

#### Immunocytochemistry

Cultures were fixed with 4% formaldehyde for 5 minutes for synaptic markers or otherwise for 10 min at room temperature (RT). Permeabilisation and blocking were performed in PBS, 10% normal serum (NS) with 0.01% Triton X-100 (NS-T) for 1hr. Cells were incubated with primary antibodies (Table S1) in 1% NS overnight at 4°C. Samples were washed with PBS and incubated in species-appropriate Alexa Fluor© secondary antibodies

(Table S2) with DAPI (1:1000) in 1% NS-T for 1 hour at RT. Images were acquired on the Opera Phenix High-content screening system, using a 40x magnification water immersion. On average, 20 images were captured per well, with 3-5 wells analysed per cell line per differentiation. Image analysis was performed using a custom pipeline developed with Harmony analysis software (v5.2, Revvity). Neurites positive for MAP2 were identified by applying a threshold of 0.4 for the region of interest. The analysis area was then expanded outwards by 5px to account for synaptic puncta, which lay slightly outside the MAP2 signal. Presynaptic (Synapsin I/II) and post-synaptic (HOMER1) puncta were detected using Method A of the 'Find Spots' function, with a threshold of >0.1 for relative intensity and a spot area restriction of ≤120 μm². Synapses were identified if HOMER1+ puncta colocalised within a 5px radius of Synapsin I/II signals.

#### Cytotoxicity BioAssay

Cytotoxicity was measured by quantifying adenylate kinase (AK) release from damaged cells using the ToxiLight™ Non-Destructive Cytotoxicity BioAssay Kit (LT07-217). Briefly, 5 µL of cell supernatant was transferred to a 384-well Greiner LUMITRAC™ plate, and 25µl of detection reagent was added to each well. After 5 minutes, luminescence was recorded using a PHERAstar® microplate reader (BMG Labtech). As a positive control, one well per cell line was treated with 10% Triton-X for 5 minutes before supernatant collection.

#### FFN206 and CypHer5E live imaging

For CypHer5E, Day 70 iPSC-DANs were treated overnight with 1:100 CypHer5E for 12 hours. Prior to imaging, cells were treated for 30 minutes with NeuroFluor™ NeuO (Stem Cell Technologies) and NucBlue (Thermo Fisher Scientific) as per manufacturers' instructions. For FFN206, iPSC-DANs were incubated for 1 hour at a concentration of 20 µm. Images were subsequently taken on an Opera Phenix High-content screening system as described above and spot properties were analysed using a custom pipeline developed with Harmony analysis software (v5.2, Revvity).

#### SynaptopHluorin live imaging

Dopamine neurons were infected with Lentivirus containing synaptopHluorin on the final replating day (Day 20). Living imaging of synaptopHluorin experiments was done using a Perkin Elmer Spinning disk confocal and Nikon eclipse Ti microscope with a Hammatsu EM-CCD camera. Cells were imaged using 500 µL of classic Tyrode's solution (124 mM NaCl, 1 mM KCl, 25 mM HEPES, 2 mM $CaCl_2$, 1 mM $MgCl_2$, 6 g/L glucose, pH 7.4), and when indicated, 500 µL of high stimulation Tyrode's with 112 mM KCl (10 mM NaCl, 112 mM KCl, 25 mM

HEPES, 4 mM $CaCl_2$, 2 mM $MgCl_2$, 6 g/L glucose, pH 7.4) was added as previously described (7), and maximum signal was measured by addition of ammonium Tyrode's (69 mM NaCl, 2.5 mM KCl, 25 mM HEPES, 2 mM $CaCl_2$, 2 mM $MgCl_2$, 6 g/L glucose, 60 mM $NH_4Cl$, pH 7.4)

#### Lentiviral production

HEK293-T LentiX (Takara Bio UK, 632180) were maintained in high glucose 1x DMEM with GlutaMAX and 10% FBS (Merck Life Science, F7524-500 mL). Prior to transfection, 15 cm dishes were coated with poly-L-ornithine for 1 hour at 37°C and dried before use. HEK cells were transfected with SynaptopHluorin construct when ~90% confluent, along with psPAX6, pMD2G and pAdVantage using Lipofectamine 3000 (Thermofisher). Twelve hours after transfection, ViralBoost (Alstem) was added with fresh media. Virus was collected two to three days after media change, harvested and spun for 10 minutes at 500g. Virus was concentrated using Lenit-X concentrator (Takara Bioscience) for 48 hours at 4°C and then spun for 1500 xg for 45 minutes at 4°C before being resuspended in 300 µL of PBS, aliquoted and stored at -80°C.

#### Statistical analysis

All statistical analyses were performed using GraphPad Prism 10.6.1 software. An outlier test using Robust regression and Outlier removal (ROUT) method (Q=1%) was performed.
